# Abnormal expression of GABA_A_ receptor sub-units and hypomotility upon loss of *gabra1* in zebrafish

**DOI:** 10.1101/2020.01.31.929455

**Authors:** Nayeli Reyes-Nava, Hung-Chun Yu, Curtis R. Coughlin, Tamim H. Shaikh, Anita M. Quintana

## Abstract

We used whole exome sequencing (WES) to determine the genetic etiology of a patient with a multi-system disorder characterized by a seizure phenotype. WES identified a heterozygous *de novo* missense mutation in the *GABRA1* gene (c.875C>T). *GABRA*1 encodes the alpha subunit of the Gamma-Aminobutyric Acid receptor A (GABA_A_R). The GABA_A_R is a ligand gated ion channel that mediates the fast inhibitory signals of the nervous system and mutations in the sub-units that compose the GABA_A_R have been previously associated with human disease. To understand the mechanisms by which *GABRA1* regulates brain development, we developed a zebrafish model of *gabra1* deficiency. *gabra1* expression is restricted to the nervous system and behavioral analysis of morpholino injected larvae suggests that the knockdown of *gabra1* results in hypoactivity and defects in the expression of other sub-units of the GABA_A_R. Expression the human GABRA1 protein in morphants partially restored the hypomotility phenotype. In contrast, the expression of the c.875C>T variant did not restore these behavioral deficits. Collectively, these results represent a functional approach to understand the mechanisms by which loss of function alleles cause disease.

## INTRODUCTION

Rare disorders affect 4-8% of the global population (Boycott et al., 2013) and approximately 80% of these disorders are predicted to have a genetic etiology (Bick et al., 2019). In recent years, whole exome sequencing (WES) emerged as a diagnostic tool for patients with rare disorders of unknown origin (Sawyer et al., 2015; Tetreault et al., 2015). The success of WES has provided a unique window of opportunity to identify disease related genes in humans and it is predicted that gene identification of rare disorders has the potential to contribute to our knowledge of other, more complex genetic disorders (Danielsson et al., 2014; Koboldt et al., 2013). Most importantly, studies of rare disorders have demonstrated that WES can be successful with very few subjects and/or using a trio based approach (Gilissen et al., 2011; Gilissen et al., 2012).

In 2013, the Undiagnosed Disease Network (UDN) was founded and includes seven clinical sites across the nation, a coordinating center, two DNA sequencing centers, a model organism screening center, a metabolomics core, and a central biorepository (Macnamara et al., 2019). Within the first 4 years of the UDN, the sequencing centers identified 956 genes associated with human disease, 375 of them have not previously been associated with disease (Wangler et al., 2017). This is a staggering number, as it suggests that nearly 1/3 of the genes identified are of unknown function. These data strongly support the need for *in vivo* functional analysis of gene function.

Here we describe the identification of a putative disease variant and perform *in vivo* functional analysis of gene function using genetic loss of function. We describe a patient who presented with a severe seizure disorder, intellectual disability, cardiac arrhythmia, and non-verbal speech. We identified a heterozygous *de novo* missense mutation in the *GABRA1* gene (c.875C>T), which resulted in a single amino acid substitution in one of the three known transmembrane domains (p.Thr292Ile). *GABRA1* is located on chromosome 5 and encodes the alpha (α) sub-unit of the multi-sub-unit gamma-aminobutyric acid receptor (GABA_A_R). The GABA_A_R is the primary inhibitory receptor of the central nervous system and the c.875C>T variant was previously associated with epileptic phenotypes in the Epi4K consortium (Epi4K Consortium et al., 2013). Although mutations in *GABRA1* have been associated with disease, the molecular and cellular mechanisms by which *GABRA1* regulates neural development are not completely understood. Consequently, we performed functional analysis in the developing zebrafish embryo.

Zebrafish are a cost-effective model organism and nearly 75% of their genome is conserved with humans (Ackermann and Paw, 2003; Reyes-Nava, Nayeli G. et al., 2018). Additionally, they are highly amenable to genetic manipulation. To ascertain the function of *GABRA1* during development and behavior, we performed morpholino mediated knockdown of the zebrafish ortholog of *GABRA1*. We analyzed the behavioral and molecular consequences associated with knockdown of *gabra1*. Morphants exhibited hypomotility as indicated by swim speed and total distance swam. This hypomotility was accompanied by distinct changes in expression of the major sub-units of the GABA_A_R, including decreased expression of the β2 and γ2 transcripts. Despite this decrease in the expression of unique GABA_A_R sub-units, morphants continued to respond to treatment with pentylenetetrazol (PTZ), a potent antagonist of the GABA_A_R, indicating that morphants continue to produce an active GABA_A_R even in the absence of adequate *gabra1* expression.

## RESULTS

### Subject

The subject initially presented to care at three months of age with infantile spasms that evolved into Lennox-Gastaut syndrome. Seizure activity included a light-sensitive myoclonic epilepsy and generalized tonic clonic seizures. The seizures were treated with adrenocorticotropic hormone (ACTH), multiple antiepileptic medications, and ketogenic diet, although seizure activity continued to occur daily.

His clinical course was also marked for hypotonia, visual impairment, developmental delay and bilateral neuromuscular hip dysplasia. He had a Torsades de pointes cardiac arrest during an acute illness, with normal cardiac function outside of the acute event. Laboratory findings included lactic acidosis (peak serum lactate 4.18, ref range 0.5-2.0 mM) and metabolic acidosis. Multiple diagnoses were suggested based on his clinical history including a channelopathy and primary mitochondrial dysfunction. In previous clinical testing, the patient was negative for mutations in a panel of genes including *ADSL, ALDH7A1, ARX, ATP6AP2, CDKL5, CLN3, CLN5, CLN6, CLN8, CNTNAP2, CTSD, FOXG1, GABRG2, GAMT, KCNQ2, KCNQ3, MECP2, MFSD8, NRXN1, PCDH19, PNKP, PNPO, POLG, PPT1, SCN1A, SCN1B, SCN2A, SLC25A22, SLC2A1, SLC9A6, SPTAN1, STXBP1, TCF4, TPP1, TSC1, TSC2, UBE3A,* and *ZEB2.* In order to investigate the underlying genetic etiology for his complex medical history, the subject and his parents were enrolled into a research protocol approved by the Colorado Multiple Institutional Review Board (COMIRB #07-0386).

### Whole-Exome Sequencing

WES was performed on a male subject and his unaffected parents to obtain over 70X coverage of targeted exons in each sample (Table S1). A large number of variants (106,737 variants) were detected in the patient after applying appropriate quality measures as described in Table S2. Our downstream analyses focused on nonsynonymous coding variants, coding InDels (insertions/deletions <50 bp), and variants affecting splice-sites as they are more likely to have a functional impact on the gene product and hence more likely to be pathogenic (9,631 variants). Common variants with minor allele frequency (MAF) greater than 1% in dbSNP137 were filtered out. Our analysis identified 1,846 rare variants in the patient and these were considered for further analysis. Parental WES data was used to detect the pathogenic variant under various inheritance models including dominant (*de novo* mutations) and recessive (compound heterozygous, homozygous, and X-linked hemizygous mutations) models. This resulted in identification of seven candidate genes (Table S2). These included *de novo* variants in *CACNA1C, GABRA1,* and compound heterozygous variants in *SCNN1B*, *FNIP1, TTN*, *OTOG*, and *FAT4.*

Additional evaluation of each candidate gene according to the criteria described in MATERIAL AND METHODS section identified 2 top priority candidate genes, including the *de novo* variant in *GABRA1* under a dominant model and compound heterozygous variants in *TTN* under a recessive model (Table S2). *TTN* encodes Titin, a sarcomeric protein involved in the assembly of cardiac and skeletal muscle. The second candidate gene, *GABRA1* has been associated with early infantile epileptic encephalopathy (EIEE19; MIM: 615744) and juvenile myoclonic epilepsy (EJM4, EJM5; MIM: 611136) and therefore, became the primary putative candidate gene based on clinical phenotype. Both parents had normal alleles but the subject had a heterozygous missense variant in *GABRA1*(NM_000806.5:c.875C>T, NP_000797.2:p.Thr292Ile) that results in a change in protein sequence (Fig. 1A). Sanger sequencing also confirmed that the variant is *de novo* (Fig. 1B) and mostly likely the result of a germline mutation. Amino acid Thr292 is highly conserved evolutionarily between multiple vertebral species (Fig. 1C) according to several conservation algorithms (PhyloP: 7.66; PhastCons: 1; GERP: 5.8). Notably, multiple mutation prediction algorithms predict this variant to be deleterious (CADD: 33; PolyPhen2 = probably damaging (1); PROVEAN = deleterious (−5.34); SIFT = damaging (0); MutationTaster: disease causing (0.99999)). Amino acids from position 279 to 300 of GABRA1 form a functionally important transmembrane helix domain (TM2) that is critical for overall functionality (Fig. 1D). The significance of p.The292Ile variant in the subject is further supported by previous studies, which have established that *de novo* mutations in the first three transmembrane domains (TM1, TM2, and TM3) are associated with neurological and epileptic conditions (Kodera et al., 2016). Most importantly, the c.875C>T heterozygous variant has been reported by the Epi4K consortium (Epi4K Consortium et al., 2013). Collectively, these data provide strong evidence that the heterozygous c.875C>T missense variant in *GABRA1* is likely pathogenic.

**Figure 1:**
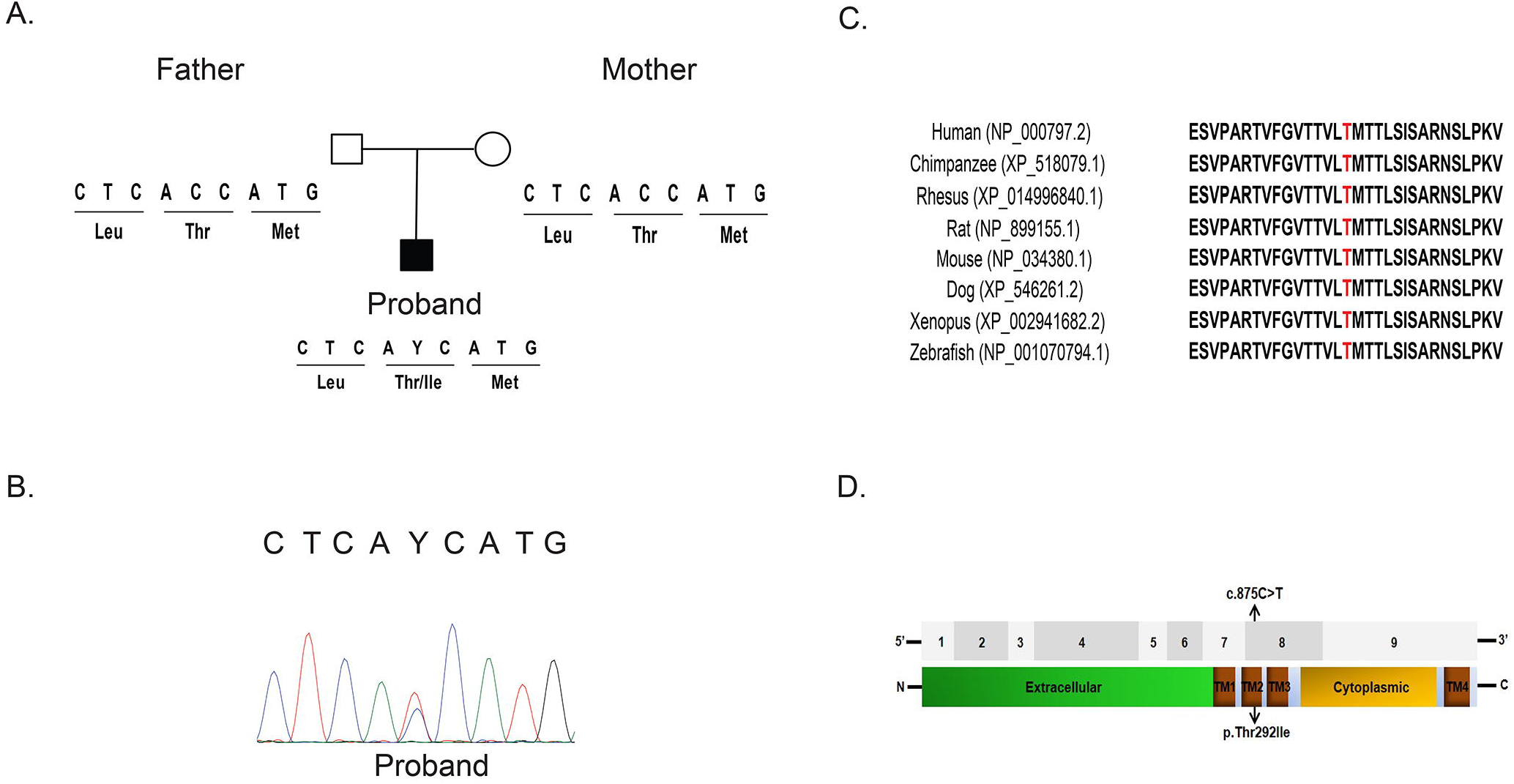
Identification of Pathogenic Variants in the *GABRA1* gene. (A) Depiction of a *de novo* missense variant c.875C>T (p.Thr292Ile) in the patient and his unaffected parents. (B) Partial chromatograms demonstrating Sanger Sequencing validation in the Proband. (C) Comparative analysis of the GABRA1 protein from multiple species. Thr292 (highlighted in red) and its neighboring amino acids are evolutionarily conserved. Protein sequences were obtained from NCBI Protein database or Ensembl. (D) Top: Annotation of the nine coding exons in the *GABRA1* gene. Bottom: The GABRA1 protein includes an extracellular domain, a cytoplasmic domain and four transmembrane domains (TM1-4) (annotated by Universal Protein Resource, UniProt). Location of variant identified in the patient is indicated by arrows within TM2.

### Expression Patterns of the zebrafish ortholog of *GABRA1*

In order to understand the mechanisms by which *GABRA1* regulates development, we used the zebrafish (*Danio rerio*) as a model organism. We first confirmed the spatial and temporal expression patterns of zebrafish *gabra1* using whole-mount in situ hybridization (WISH). We performed WISH at 1, 2, and 3 days post fertilization (DPF). *gabra1* expression was localized to the developing nervous system at each time points with the broadest expression at 1 DPF (Fig. 2). Over the course of development the expression of *gabra1* became more restricted to the midbrain-hindbrain regions (Fig. 2) consistent with previously published work (Monesson-Olson et al., 2018; Samarut et al., 2018).

**Figure 2:**
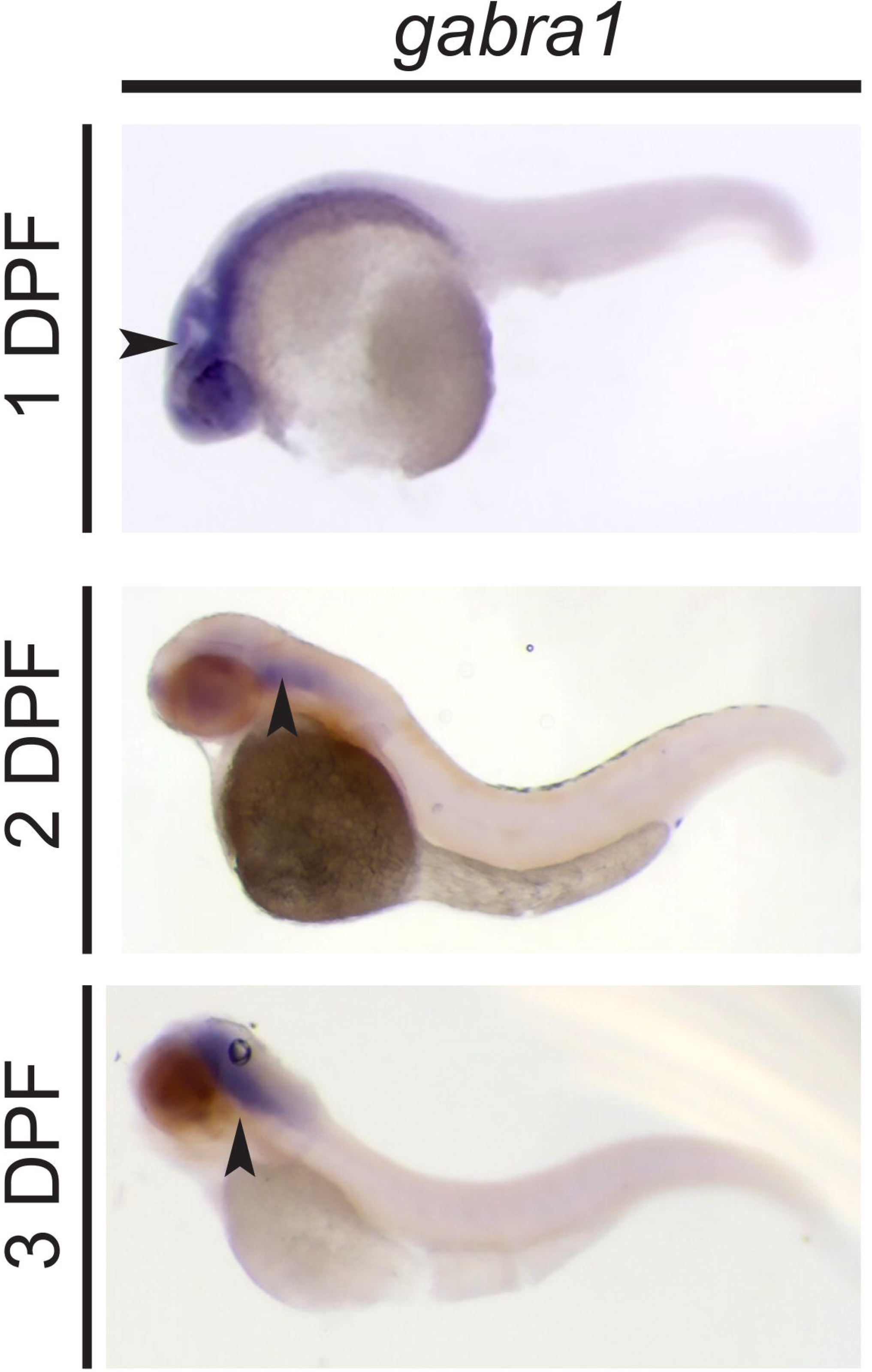
*gabra1* expression in the developing zebrafish. Whole mount *in situ* hybridization (WISH) was performed at 1, 2, 3 days post fertilization (DPF) with an anti-sense *gabra1* probe. Arrows indicated the expression of *gabra1* at each developmental stage.

### Gabra1 regulates zebrafish larval motility

Mutations in *GABRA1* have been associated with epileptic phenotypes (Cossette et al., 2002; Maljevic Snezana et al., 2006; Lachance-Touchette et al., 2011; Kodera et al., 2016; Farnaes et al., 2017; Nolan and Fink, 2018) and behavioral assays to monitor seizure like behaviors in zebrafish have emerged (Baraban et al., 2005; Reyes-Nava, Nayeli G. et al., 2018). Consequently, we developed a protocol using the Zebrabox behavioral unit to monitor swim speed and total distance swam in larvae injected with anti-sense morpholinos that inhibit either the translation of *gabra1* or mRNA splicing. Embryos were injected at the single cell stage with randomized control morpholinos (RC), translational targeting morpholinos (tbMO), or mRNA disrupting morpholinos (sMO) and raised to 5 DPF. Larvae were monitored according to the protocol described in the MATERIALS AND METHODS for swim speed and total distance swam. As shown in Figure 3, the tbMO was associated with a statistically significant (p<0.001) reduction in total swim speed (Fig. 3A) and decreased total distance swam (Fig. 3B) consistent with a hypomotility phenotype. The decrease in speed and distance was observed in both light and dark conditions at 5 DPF (Fig. 3C). Importantly, injection of an equivalent concentration of sMO induced a hypomotility phenotype (Fig. S1B,C; p=0.0263). These results are consistent with the phenotype present in morphants injected with the translational blocking morpholino. Importantly, we validated the effects of injection of sMO on mRNA splicing, which demonstrated a near 50% reduction in wildtype *gabra1* (Fig. S1A).

**Figure 3:**
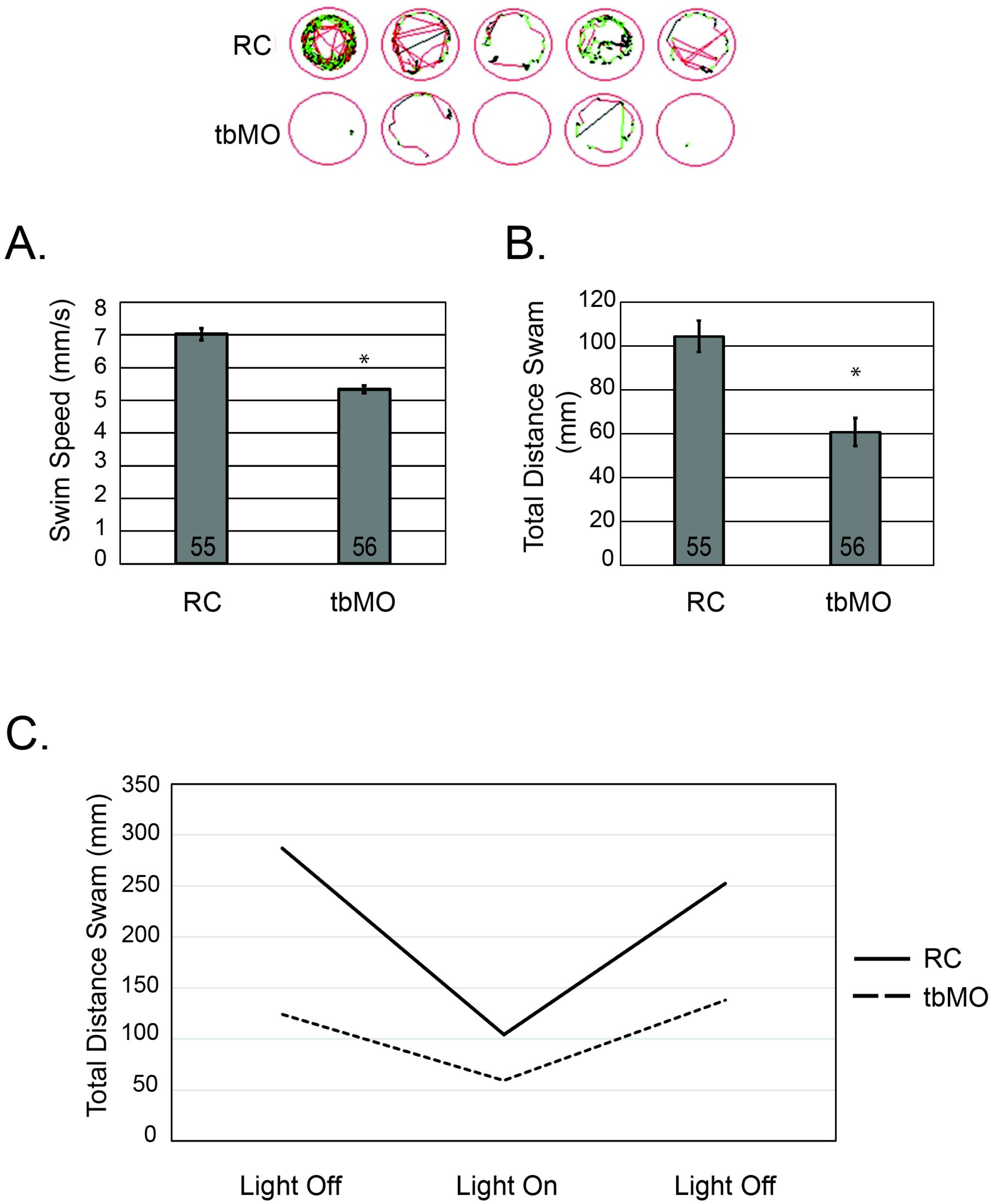
Knockdown of *gabra1* causes hypomotility. (A) Total swim speed of larvae injected with random control morpholinos (RC) or translational targeting *gabra1* morpholinos (tbMO) was determined using Zebrabox technology at 5 days post fertilization (DPF). Total number of embryos analyzed per group is depicted in the graph. *p<0.001. (B) The total distance swam was assessed at 5 DPF using Zebrabox technology. *p<0.001. Representative images of larval swim patterns are depicted above panel (A) and (B). (C) The total distance swam was assessed in *gabra1* morphants and random control injected larvae at 5 DPF. Distance was calculated without light, after the onset of light for a 5 minute duration, and an additional period without light.

Next, we sought to restore the hypomotility phenotype in morphants (tbMO) by co-injection of *GABRA1* encoding mRNA. Embryos were injected at the single cell stage with RC morpholinos, tbMO morpholinos, *GABRA1* mRNA, or a combination of *GABRA1* mRNA with RC or tbMO morpholinos. Injection of the tbMO caused a statistically significant decrease in the total distance swam (p= 0.000161) and the overall swim speed (p=0.036384) (Fig.4A,B; RC relative to tbMO). Injection of *GABRA1* encoding mRNA had no significant effect on speed or distance at a concentration of 1000pg/embryo (Fig. 4A,B; mRNA and RC+). The co-injection of the tbMO and *GABRA1* encoding mRNA at 1000pg/embryo restored the total distance swam to normal levels (p= 0.003010689), but was not sufficient to restore the deficits in overall speed to control levels (Fig. 4A,B). Thus, co-injection of 1000pg of *GABRA1* encoding mRNA with the tbMO produced a partial rescue of the observed phenotype. Injection of *GABRA1* mRNA at higher concentrations was accompanied by some degree of toxicity (cardiac edema and death) and therefore, additional rescue experiments with higher concentrations could not be attempted.

**Figure 4:**
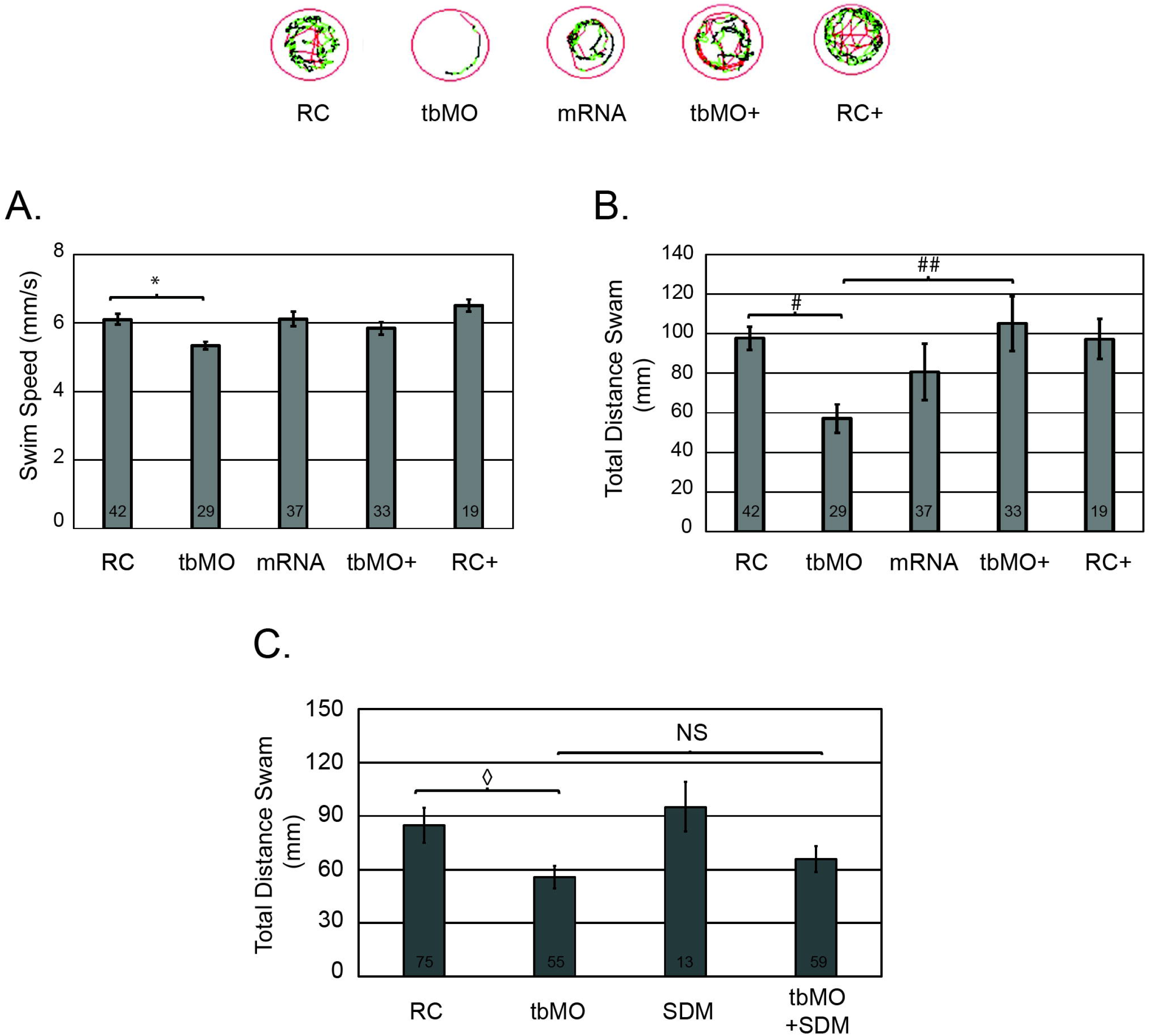
Restoration of hypomotility by co-injection of human mRNA variants. (A) Total swim speed of larvae injected with random control morpholinos (RC), translational targeting *gabra1* morpholinos (tbMO), *GABRA1* encoding mRNA (1000pg/embryo), RC with *GABRA1* mRNA (RC+), or tbMO with *GABRA1* mRNA (tbMO+) was determined using Zebrabox technology at 5 days post fertilization (DPF). Total number of embryos analyzed per group is depicted in the graph. (B) The total distance swam was assessed at 5 DPF using Zebrabox technology for each of the conditions in (A). *p=0.036384 and ^#^p=0.000161 and ^##^p=0.003010689. Representative images of larval swim patterns are depicted above panel (A) and (B). (C) Total distance swam of larvae injected with RC, tbMO, GABRA1 c.875C>T (SDM) encoding mRNA, or tbMO with GABRA1 c.875C>T encoding mRNA (tbMO+SDM) was determined at 5 DPF. Total number of animals is indicated in the graph. ^◊^ p=0.0153.

### The c.875C>T *GABRA1* variant does not restore the hypomotility phenotype in morphants

The functional consequences of the c.875C>T variant are currently unknown. Therefore, we asked whether expression of the c.875C>T variant was sufficient to restore the hypomotility induced by knockdown of *gabra1*. Embryos were injected at the single cell stage with RC morpholinos, tbMO morpholinos, *GABRA1* c.875C>T mRNA (SDM), or a combination of *GABRA1* mRNA with tbMO morpholinos. Consistent with previous experiments, injection of the tbMO morpholino caused a significant reduction (p=0.0153) in the total distance swam relative to embryos injected with the RC (Fig. 4C). Interestingly, the co-injection of the mRNA encoding the c.875C>T (SDM) and the tbMO was unable to restore the total distance traveled to control levels (Fig. 4C). Importantly, the injection of the *GABRA1* c.875C>T variant (SDM) at 1000pg/embryo had no significant effects on the total distance swam (Fig. 4C).

### The expression of *gabrb2* and *gabrg2* are decreased in *gabra1* morphants

Previous studies suggest that approximately 60% of all GABA_A_Rs consist of two α1, two β2, and one γ2 subunits (Sigel and Steinmann, 2012). We hypothesized that the knockdown of *gabra1*, which encodes the α1 sub-unit would alter the sub-unit composition of the GABA_A_R. To begin to test this, we analyzed the expression of the genes that encode the β2 and γ2 sub-units. As shown in Figure 5A, knockdown of *gabra1* caused a decrease in the expression of *gabrb2* (β2) and *gabrg2* (γ2). We next measured the expression of other alpha sub-units in *gabra1* morphants. As shown in Figure 5A, injection of the tbMO was associated with increased expression of *gabra6a* and *gabra6b*, but only gabra6b was statistically significant across biological triplicates. A similar expression pattern of *gabra6a* and *gabra6b* was observed upon injection of the sMO (Fig. S1D), with both genes demonstrating a statistically significant increase in expression. We did not detect a statistical change in the expression of any other α sub-unit across either the tbMO or the sMO (Figs 5A, S1D).

**Figure 5:**
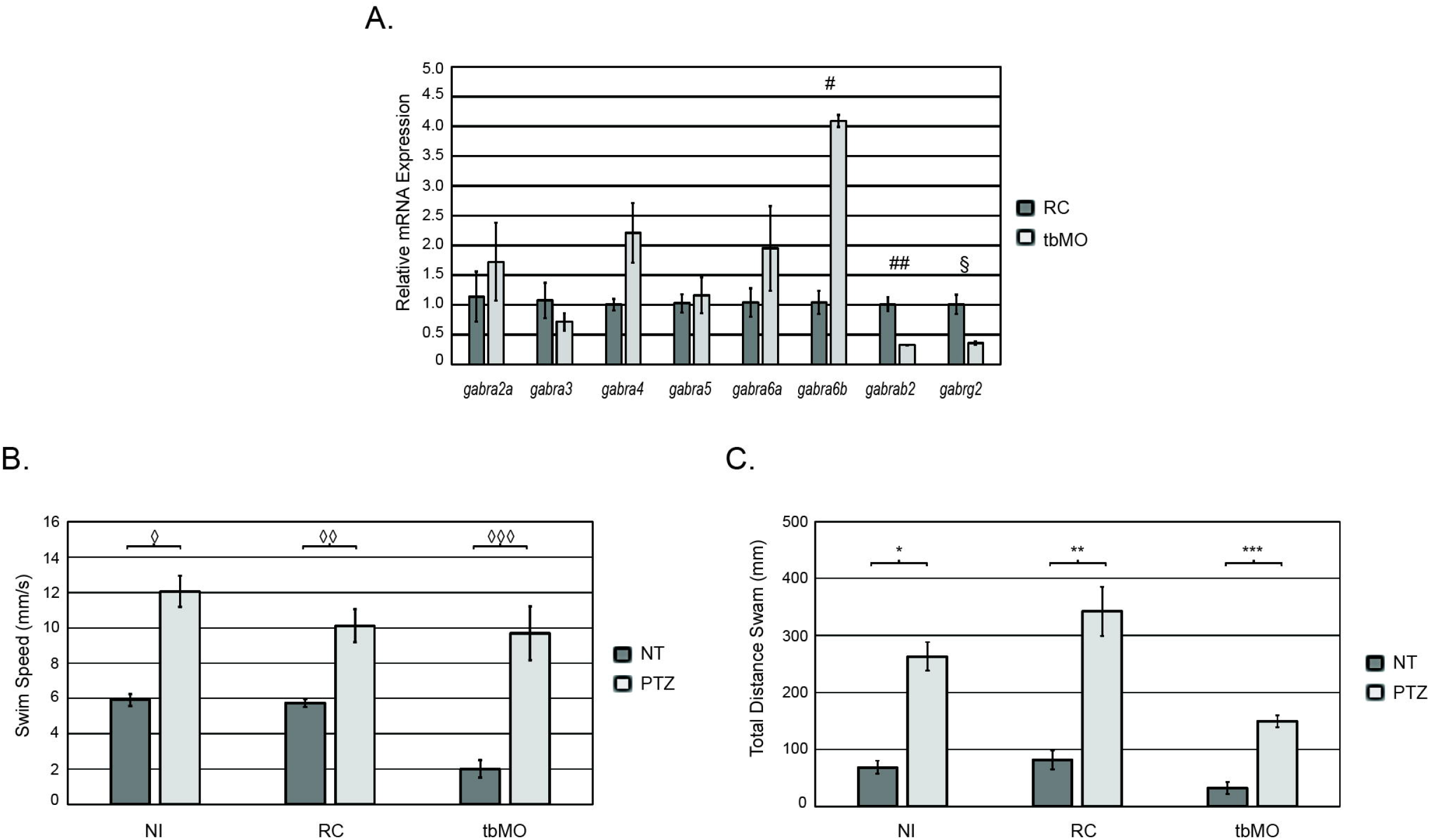
Molecular and behavioral responses of *gabra1* morphants. (A) Quantitative real time PCR (QPCR) was performed at 5 days post fertilization (DPF) to measure the expression of each gene indicated. Total RNA was isolated from random control injected embryos (RC) or translational blocking *gabra1* morpholinos (tbMO). Error bars represent standard deviation. Expression was measured in biological triplicate. ^#^p=0.0016, ^##^p=0.003681, ^§^p= 0.009. (B) Total swim speed was assessed at 5 DPF in non-injected larvae, larvae injected with RC, or larvae injected with MO treated with 10uM pentylenetetrazole (PTZ). ^◊^p=6.74E-06, ^◊◊^ p=5.34E-07, ^◊◊◊^p=1.019E-05. (C) Total distance travelled was assessed at 5 DPF in each of the groups described in (B). *2.14E-05. **p=3.92141E-05, ***p=0.0002636.

We sought to build upon these data by determining whether morphant larvae had an intact receptor capable of responding to pentylenetetrazol (PTZ), an antagonist of the GABA_A_R. Non-injected wildtype embryos treated with 10mM PTZ exhibit short convulsions and a whirlpool swimming behavior with a 2-fold increase in swim speed (p= 6.74E-06) and an approximate 6-fold increase in total distance swam (p= 2.14E-05) (Fig. 5B,C). These phenotypes were consistently observed in larvae injected with RC morpholinos as the RC morpholino had no affect on larval behavior or their response to PTZ. Interestingly, *gabra1* morphants (tbMO) responded normally to PTZ according to both distance and speed measurements (Fig. 5B,C). Collectively, these data demonstrate that knockdown of *gabra1* alters the expression of unique GABA_A_R subunits, although, morphants continue to respond to PTZ treatment.

## DISCUSSION

We have identified an individual presenting with multi-system disorder carrying a *de novo* missense variant in the *GABRA1* gene (NM_000806.5:c.875C>T, NP_000797.2:p.Thr292Ile). The *GABRA1* gene encodes the α1 sub-unit of the GABA_A_R, which mediates the fast inhibitory synapses of the nervous system. GABA_A_Rs are pentameric and can be composed of different combinations of the following components: six α subunits, three β subunits, three γ subunits, three ρ subunits, one ε, δ, θ, or π subunits. Of these sub-units, mutations in *GABRA1* (Cossette et al., 2002, 1; Kodera et al., 2016, 1; Macdonald and Gallagher, 2015; von Deimling et al., 2017), *GABRA6* (Hernandez et al., 2011), *GABRB2* (Macdonald and Gallagher, 2015), *GABRB3* (DeLorey et al., 1998; von Deimling et al., 2017), *GABRG2* (von Deimling et al., 2017), and *GABRD* (Macdonald and Gallagher, 2015) have been associated with epileptic phenotypes (reviewed in (Hirose, 2014)). Most importantly, in a recent international collaboration (Epi4K Consortium), the heterozygous *de novo* p.Thr292Ile variant we describe here was identified in a male patient diagnosed with infantile spasms (Epi4K Consortium et al., 2013). The individual studied in the Epi4K study had febrile seizures at the age of 1-month and at 15-months of age, his electroencephalogram (EEG) showed bursts of generalized spike and wave (GSW) at 2.5 Hz with multiple foci of epileptiform activity. He presented with features of generalized tonic clonic (GTC) and myoclonic seizures. He was developmentally delayed, hypotonic, and did not speak at 18-months of age with additional features that include esotropia, poor vision, abnormal electroretinogram, and a head circumference at 5th percentile. The subject reported here was diagnosed with seizure disorder, intellectual disability, vision loss, and was non-verbal; phenotypes consistent with the previously identified case. Additionally, the p.Thr292Ile variant is present in one of the 3 transmembrane domains of the GABRA1 protein and these domains have been associated with epileptic phenotypes (Kodera et al., 2016). Collectively, these data strongly suggest that the heterozygous mutation p.Thr292Ile causes a complex disorder characterized by a severe seizures. This is supported by the fact that there are at least two subjects with overlapping phenotypes harboring this variant.

It is not yet known how mutations in the GABRA1 transmembrane domain result in seizure like phenotypes. Genetic knockout mice have been developed to understand how mutations in *Gabra1* (mouse) affect GABA_A_R function, but the results have been difficult to interpret, as the deletion of *Gabra1* (mouse) causes strain and sex specific phenotypes (Arain et al., 2012). Due to these strain differences, additional systems have been developed including a zebrafish harboring a mutation in the *gabra1* gene. Interestingly, mouse models of *Gabra1* deletion are viable, but the homozygous deletion of *gabra1* in fish is lethal (Samarut et al., 2018). Despite this lethality, mutant zebrafish survive to several weeks post fertilization, which has allowed for the characterization of *gabra1* function in fish at 7-10 weeks post fertilization (Samarut et al., 2018).

In this report, we demonstrate that morpholino mediated knockdown of *gabra1* in zebrafish leads to hypomotility in the presence and absence of light. We perform our studies at 5 DPF, during the larval stage, prior to the onset of feeding or sexual dimorphism, but after swim bladder formation. *gabra1* morphants consistently demonstrated with reduced swim speed and reduced overall distance travelled relative to control. These data are consistent with Samarut et. al., who demonstrated that mutation of *gabra1* results in hypomotility, albeit at a later stage in development that would be equivalent to a juvenile onset (Samarut et al., 2018). In contrast to Samarut et. al., we did not observe overt indications of myoclonic seizures at any time point in our protocol. For example, within the first minute of light exposure, Samarut and colleagues observed intense seizures characterized by convulsions, uncontrolled movements, and whirlpool swim behavior. This phenotype was not observed in morphant animals (data not shown). This can likely be attributed to the fact that our study is performed using a knockdown of *gabra1*, which maybe more consistent with the heterozygous phenotypes reported by Samarut and colleagues at 4 DPF. To address the function of the c.875C>T *GABRA1* variant, we performed restoration experiments in which this variant was co-injected with *gabra1* targeting morpholinos. Co-injection of mRNA encoding the c.875C>T variant did not restore the hypomotility phenotype present in morphants, whereas co-injection of wildtype *GABRA1* restored the total distance travels to control levels. These data suggest that the c.875C>T variant is a loss of function allele, however, future studies characterizing the function of this variant are warranted. Should this allele be a loss of function allele, morpholino mediated knockdown is an alternative approach towards understanding the mechanisms by which the c.875C>T allele causes disease.

We further demonstrate that knockdown of *gabra1* causes abnormal expression of other sub-units of the GABA_A_R. Despite changes in the expression of various GABA_A_R sub-units, morphant animals continue to respond to PTZ stimulus. PTZ is a potent antagonist of the GABA_A_R and treatment of wildtype larvae with PTZ induces a myoclonic seizure (Afrikanova et al., 2013; Baraban et al., 2005) because PTZ binds directly to the GABA_A_R resulting in disinhibition (Huang et al., 2001). The continued response of morphants to PTZ, suggests that these embryos maintain the ability to produce some form of the GABA_A_R. Consistent with this hypothesis, we observed increased expression of *gabra6a and gabra6b* mRNA, which encode the two zebrafish α6 sub-units of the GABA_A_R. Other sub-units did not demonstrate consistent changes in expression across multiple morpholinos or biological replicates. Interestingly, mutations in *GABRA6*, which encodes the α6 sub-unit are associated with disease (Hernandez et al., 2011). Thus, it is unclear whether the hypomotility phenotype observed is the direct result of a lack of g*abra1* or the up-regulation of *gabra6*. Future studies analyzing the function of α6 and other alpha sub-units in *gabra1* mutant animals are needed.

The gene expression changes we observe are strongly supported by previous conclusions in mice with mutations in the *Gabra1* gene (Arain et al., 2015; Zhou et al., 2015). Recent work in zebrafish has demonstrated that the homozygous nonsense mutation of *gabra1*, does not disrupt overall brain structure or the total number of GABAergic cells, but does influence the brain transcriptome (Samarut et al., 2018). Collectively, these data raise the possibility that other alpha sub-units may compensate for the loss of *gabra1,* ultimately producing unique compositions of the GABA_A_R. The function of these receptors is unknown. But it is conceivable that the production of GABA_A_R with unique sub-unit composition in incorrect regions of the brain might underlie the impaired synapse formation observed in zebrafish harboring germline mutations in the *gabra1* gene (Samarut et al., 2018).

We provide strong evidence that heterozygous *de novo* mutation of *GABRA1* is associated with a multi-system disorder characterized by severe seizures. We further characterized the developmental and behavioral defects associated with knockdown of *gabra1* in zebrafish. Behaviorally, morphant animals present with hypomotility at 5 DPF measured by reduced swim speed and total distance traveled. These deficits coincide with significant changes in the expression of GABA_A_R sub-units and cannot be restored by the *de novo* c.875C>T allele. Although a zebrafish harboring a mutation in the *gabra1* gene has recently been created, detailed behavioral analysis was performed at the juvenile stage (weeks post fertilization). Here we complement previous studies using a morpholino mediated knockdown approach, as the homozygous deletion of *gabra1* was lethal (Samarut et al., 2018). Our behavioral study is the first to our knowledge that comprehensively characterizes the phenotype of *gabra1* deletion during early development (DPF as opposed to weeks post fertilization). We observed hypomotility consistent with previous studies in zebrafish and our study likely informs about specific types of mutations, those of which result in loss of function alleles. Importantly, our restoration experiments with the c.875C>T allele suggest that this allele is in fact a loss of function allele. Thus, morpholino mediated studies might provide insight into the mechanisms by which loss of function alleles cause disease.

## MATERIALS AND METHODS

### Animal Husbandry

For all experiments, embryos were obtained by crossing AB wildtype or Tupfel Long Fin wildtype. Fish were maintained at The University of Texas at El Paso according to the Institutional Animal Care and Use committee (IACUC) guidelines (Protocol Number 811689-5). They were maintained and bred in groups of two females and two to four males. The collected zebrafish embryos were kept in egg water consisting of 0.03% Instant Ocean (Aquaneering, San Diego, CA) in D.I. water at 28°C.

### Whole-Exome Sequencing and Data Analysis

High quality, unamplified, and unfragmented genomic DNA (A260/A280 ≥ 1.8 and A260/A230 ≥ 1.9) was extracted from whole blood obtained from the subject and his parents using the Puregene Blood kit from Qiagen (Valencia, CA). Whole exome sequencing was performed using the service provided by Beijing Genomics Institute (Cambridge, MA). Details of data analysis were similar to the procedure as previously described (Epi4K Consortium et al., 2013). Approximately 78 to 168 million, 100 bp, paired-end reads (>70X) were obtained and mapped to the reference human genome (GRCh37/hg19) using Burrows-Wheeler Aligner (BWA) (Li and Durbin, 2009, 200) (summarized in Table S1). Variants were determined by the utilities in the SAMtools (Li et al., 2009) and further annotated with SeattleSeq. Filtering and the test of inheritance model was performed using tools available in Galaxy (Goecks et al., 2010). Variants were filtered against dbSNP build 137, 1000 Genomes (November 23, 2010 release version), Exome Variant Server (EVS, ESP6500SI-V2) and Exome Aggregation Consortium ExAC browser (version 0.3). Rare variants were identified as a variant with a minor allele frequency (MAF) less than 1% using dbSNP137. The sequence data from the family was then used to test for causal variants under different inheritance models, including dominant (*de novo* mutations) and recessive (compound heterozygous, homozygous, and X-linked hemizygous mutations) models. In the dominant model, variants found in any database (dbSNP, 1000 Genomes, EVS, ExAC) were removed from the top candidacy list. In the recessive model, autosomal variants which had homozygotes found in the databases, such as EVS and ExAC, (or variants on chrX or chrY with hemizygotes in databases) were deleted from the top candidacy list.

### Sanger Sequencing Verification

Sanger sequencing was used to validate the variant described. Briefly, primers were used to amplify the PCR product (fwd 5’-GCTGTFATAGGGTGGAGGTG-3’, rev 5’GCTATCAACGCCATTGTGAA-3’) using 1X GoTaq Green (Promega, Madison, WI) with a final primer concentration of 0.2uM. Reaction parameters for PCR include an initial cycle at 95°C for 10 minutes, followed by 30 cycles of 95°C for 30 seconds, 60°C for 30 second, and 72°C for 1 minute, finishing with extension at 72°C for 5 minutes. Amplified PCR products were sequenced using the PCR primers as sequencing primers. Variations detected in *GABRA1* were assigned using cDNA accession number NM_000806.5.

### Whole mount *in situ* hybridization and Injections

WISH was performed as previously described (Thisse and Thisse, 2008). Embryos were harvested at 1, 2, and 3 DPF and fixed in in 4% paraformaldehyde (PFA) (Electron Microscopy Sciences, PA) for 1 hour at room temperature (RT). Embryos were dehydrated using a methanol: PBS gradient and stored in 100% methanol overnight in −20°C. Embryos were rehydrated using PBS: Methanol gradient, washed in PBS with 0.1% Tween 20 and permeabilized with proteinase K (10ug/ml) for the time indicated by Thisse and Thisse (Thisse and Thisse, 2008). Permeabilized embryos were prehybridized in hybridization buffer (HB) (50% deionized formamide (Fisher, Waltham, MA), 5X SSC (Fisher, Waltham, MA), 0.1% Tween 20 (Fisher, Waltham, MA), 50μg ml^−1^ heparin (Sigma, St. Louis, MO), 500μg ml^−1^ of RNase-free tRNA (Sigma, St. Louis, MO), 1M citric acid (Fisher, Waltham, MA) (460μl for 50ml of HB) for 2-4 hours and then incubated overnight in fresh HB with probe (*gabra1* 100ng) at 70°C. Samples were washed according to protocol, blocked in 2% sheep serum (Sigma, St. Louis, MO), 2 mg ml^−1^ bovine serum albumin (BSA) (Sigma, St. Louis, MO) for 2-4 hours at RT, and incubated with anti-DIG Fab fragments (1:10,000) (Sigma, St. Louis, MO) overnight at 4°C. Samples were developed with BM purple AP substrate (Sigma, St. Louis, MO) and images were collected with a Zeiss Discovery Stereo Microscope fitted with Zen Software. The *gabra1* probe was created using primers specific to the endogenous cDNA sequence (*gabra1* ISH fwd 5’- TAAGCTGCGCTCTTCTCCTC-3’, *gabra1* ISH rev 5’-GCAGAGTCCCTTCCTCTGTG-3’).

For morpholino injections, a translational blocking morpholino (tbMO) (TCTTCCACCCCACATCATTCTCCGA) and a splice site inhibiting morpholino (sMO) (ACACGCTCTGTTGAAGCAAGAAATT) targeting *gabra1* were designed. The efficiency of knockdown for the sMO was performed with primers flanking the target site (Fwd: GACAGCCTCCTCGATGGTTA and Rev: GCAGAGTCCCTTCCTCTGTG). Each morpholino was injected independently at the single cell stage at a concentration of 1.6 ng/embryo. An equivalent concentration of randomized control morpholinos (25-N) was injected as a control. Final concentration of morpholino was determined empirically after an injection gradient was performed to determine optimal survival. For rescue experiments, the human *GABRA1* complete open reading frame was purchased from TransOMIC Technologies (Huntsville, AL). The c.875C>T *GABRA1* variant was created from the original vector obtained from TransOMIC Technologies using the QuikChange II Site-Directed Mutagenesis Kit (Fisher, Waltham, MA) with forward (TAACAACTGTGCTCATCATGACAACATTGAG) and reverse primers (GAGTTACAACAGTACGACTCGTGTCAACAAT). *In vitro* RNA was synthesized using the mMessage Machine kit (Fisher, Waltham, MA). The synthesized mRNA was injected at the single cell stage alone or in conjunction with tbMO at the indicated concentrations in the figure legends.

### Quantitative Real Time PCR (QPCR)

Total RNA was isolated from brain homogenates obtained from embryos injected with random control morpholinos or tbMO at 5 DPF using Trizol (Fisher, Waltham, MA). Reverse transcription was performed using the Verso cDNA Synthesis Kit (Fisher, Waltham, MA) and total RNA was normalized by concentration (ng) across all samples. PCR was performed in technical triplicates for each sample using an Applied Biosystems StepOne Plus machine with Applied Biosystems associated software. Sybr green (Fisher, Waltham, MA) based primer pairs for each gene analyzed are as follows: *gabra2a* fwd GATGGCTACGACAACAGGCT,*gabra2a* rev TGTCCATCGCTGTCGGAAAA, *gabra3* fwd GCTGAAGTTCGGGAGCTATG, *gabra3* rev GGAGCTGATGGTCTCTTTGC,*gabra4* fwd GACTGCGATGTACCCCACTT,*gabra4* rev ATCCAGGTCGGAGTCTGTTG, *gabra5* fwd CATGACAACACCCAACAAGC, *gabra5* rev CAGGGCCTTTTGTCCATTTA,*gabra6a* fwd TCGCGTACCCATCTTTCTTC, *gabra6a* rev CCCTGAGCTTTTCCAGAGTG,*gabra6b* fwd CGGAGGAGTGCTGAAGAAAC, *gabra6b* rev GGGAAAAGGATGCGTGAGTA, *gabrb2* fwd CCCGACACCTATTTCCTCAA,*gabrb2* rev TCTCGATCTCCAGTGTGCAG, *gabrg2* fwd ACACCCAATAGGATGCTTCG,*gabrg2* rev AGCTGCGCTTCCACTTGTAT. Analysis performed using 2^ΔΔct^. Statistical analysis of mRNA expression was performed using a T-test. All QPCR was performed in biological duplicate or triplicate using a pool of embryos (30-40) per time point.

### Behavioral Analysis and Pentylenetetrazol treatment

Embryos injected with random control morpholinos, tbMO, sMO, *GABRA1* mRNA, *GABRA1* (c.875C>T), or a combination as indicated in the figure legends were raised to 5 DPF. Behavioral analysis was performed using the Zebrabox (ViewPoint Behavior Technology, Montréal, Canada). Larvae were individually tracked for swim speed and total distance swam in a 96 well plate. The behavioral protocol (adapted from (Afrikanova et al., 2013)) was a total of 15 minutes divided into 5 minute intervals of dark/light/dark conditions. All larvae were acclimated to the dish and housing conditions for 1 hour prior to analysis. Settings for the program include a threshold of 16 and integration period of 300 seconds. Data was measured as total distance traveled (mm) and total swim speed (mm/sec) (Swim Speed= {Total distance traveled in large and small movements) (Smldist+Lardist)}/{Total duration spent by the animal in small and large movements(smldur+lardur)}. Statistical significance was determined according to a T-test. All experiments were performed in biological triplicate. For PTZ treatment, PTZ (10mM) was added directly to the 96 well plate following acclimation period. Final concentration of PTZ was determined from previously published results (Baraban et al., 2005; Afrikanova et al., 2013; Peng et al., 2016; Jin et al., 2018).

## ACKNOWLEDGEMENTS

These data could not have been completed without the patients involved in the study and without the support of Dr. Douglas Watts.

## DATA AVAILABILITY

All reagents are available upon request from the corresponding author.

## AUTHOR CONTRIBUTIONS

NR synthesized hypothesis, performing behavioral and molecular experiments, *in situ* hybridization, and all morpholino injections. HY performed bioinformatics analysis and wrote portions of the manuscript. THS and CC supervised and managed patient IRB, genetic counseling, contributed to writing the manuscript, and provided patient assessment expertise. AMQ synthesized the hypothesis, performed data management and analysis, and wrote the manuscript.

## REFERENCES

1. Ackermann, G. E. and Paw, B. H. (2003). Zebrafish: a genetic model for vertebrate organogenesis and human disorders. Front. Biosci. J. Virtual Libr. 8, d1227–1253.

2. Afrikanova, T., Serruys, A.-S. K., Buenafe, O. E. M., Clinckers, R., Smolders, I., Witte, P. A. M. de, Crawford, A. D. and Esguerra, C. V. (2013). Validation of the Zebrafish Pentylenetetrazol Seizure Model: Locomotor versus Electrographic Responses to Antiepileptic Drugs. PLOS ONE 8, e54166.

3. Arain, F. M., Boyd, K. L. and Gallagher, M. J. (2012). Decreased viability and absence-like epilepsy in mice lacking or deficient in the GABAA receptor ? 1 subunit. Epilepsia 53, e161–165.

4. Arain, F., Zhou, C., Ding, L., Zaidi, S. and Gallagher, M. J. (2015). The Developmental Evolution of the Seizure Phenotype and Cortical Inhibition in Mouse Models of Juvenile Myoclonic Epilepsy. Neurobiol. Dis. 82, 164–175.

5. Baraban, S. C., Taylor, M. R., Castro, P. A. and Baier, H. (2005). Pentylenetetrazole induced changes in zebrafish behavior, neural activity and c-fos expression. Neuroscience. 131, 759–768.

6. Bick, D., Jones, M., Taylor, S. L., Taft, R. J. and Belmont, J. (2019). Case for genome sequencing in infants and children with rare, undiagnosed or genetic diseases. J. Med. Genet.

7. Boycott, K. M., Vanstone, M. R., Bulman, D. E. and MacKenzie, A. E. (2013). Rare-disease genetics in the era of next-generation sequencing: discovery to translation. Nat. Rev. Genet. 14, 681–691.

8. Cossette, P., Liu, L., Brisebois, K., Dong, H., Lortie, A., Vanasse, M., Saint-Hilaire, J.-M., Carmant, L., Verner, A., Lu, W.-Y., et al. (2002). Mutation of GABRA1 in an autosomal dominant form of juvenile myoclonic epilepsy. Nat. Genet. 31, 184–189.

9. Danielsson, K., Mun, L. J., Lordemann, A., Mao, J. and Lin, C.-H. J. (2014). Next-generation sequencing applied to rare diseases genomics. Expert Rev. Mol. Diagn. 14, 469–487.

10. DeLorey, T. M., Handforth, A., Anagnostaras, S. G., Homanics, G. E., Minassian, B. A., Asatourian, A., Fanselow, M. S., Delgado-Escueta, A., Ellison, G. D. and Olsen, R. W. (1998). Mice lacking the beta3 subunit of the GABAA receptor have the epilepsy phenotype and many of the behavioral characteristics of Angelman syndrome. J. Neurosci. Off. J. Soc. Neurosci. 18, 8505–8514.

11. Epi4K Consortium, Epilepsy Phenome/Genome Project, Allen, A. S., Berkovic, S. F., Cossette, P., Delanty, N., Dlugos, D., Eichler, E. E., Epstein, M. P., Glauser, T., et al. (2013). De novo mutations in epileptic encephalopathies. Nature. 501, 217–221.

12. Farnaes, L., Nahas, S. A., Chowdhury, S., Nelson, J., Batalov, S., Dimmock, D. M., Kingsmore, S. F. and RCIGM Investigators (2017). Rapid whole-genome sequencing identifies a novel GABRA1 variant associated with West syndrome. Cold Spring Harb. Mol. Case Stud. 3,.

13. Gilissen, C., Hoischen, A., Brunner, H. G. and Veltman, J. A. (2011). Unlocking Mendelian disease using exome sequencing. Genome Biol. 12, 228.

14. Gilissen, C., Hoischen, A., Brunner, H. G. and Veltman, J. A. (2012). Disease gene identification strategies for exome sequencing. Eur. J. Hum. Genet. EJHG. 20, 490–497.

15. Goecks, J., Nekrutenko, A., Taylor, J. and The Galaxy Team (2010). Galaxy: a comprehensive approach for supporting accessible, reproducible, and transparent computational research in the life sciences. Genome Biol. 11, R86.

16. Hernandez, C. C., Gurba, K. N., Hu, N. and Macdonald, R. L. (2011). The GABRA6 mutation, R46W, associated with childhood absence epilepsy, alters 6β22 and 6β2 GABA(A) receptor channel gating and expression. J. Physiol. 589, 5857–5878.

17. Hirose, S. (2014). Mutant GABA(A) receptor subunits in genetic (idiopathic) epilepsy. Prog. Brain Res. 213, 55–85.

18. Huang, R.-Q., Bell-Horner, C. L., Dibas, M. I., Covey, D. F., Drewe, J. A. and Dillon, G. H. (2001). Pentylenetetrazole-Induced Inhibition of Recombinant γ-Aminobutyric Acid Type A (GABAA) Receptors: Mechanism and Site of Action. J. Pharmacol. Exp. Ther. 298, 986–995.

19. Jin, M., He, Q., Zhang, S., Cui, Y., Han, L. and Liu, K. (2018). Gastrodin Suppresses Pentylenetetrazole-Induced Seizures Progression by Modulating Oxidative Stress in Zebrafish. Neurochem. Res. 1–14.

20. Koboldt, D. C., Steinberg, K. M., Larson, D. E., Wilson, R. K. and Mardis, E. R. (2013). The next-generation sequencing revolution and its impact on genomics. Cell. 155, 27–38.

21. Kodera, H., Ohba, C., Kato, M., Maeda, T., Araki, K., Tajima, D., Matsuo, M., Hino-Fukuyo, N., Kohashi, K., Ishiyama, A., et al. (2016). De novo GABRA1 mutations in Ohtahara and West syndromes. Epilepsia. 57, 566–573.

22. Lachance-Touchette, P., Brown, P., Meloche, C., Kinirons, P., Lapointe, L., Lacasse, H., Lortie, A., Carmant, L., Bedford, F., Bowie, D., et al. (2011). Novel α1 and γ2 GABAA receptor subunit mutations in families with idiopathic generalized epilepsy. Eur. J. Neurosci. 34, 237–249.

23. Li, H. and Durbin, R. (2009). Fast and accurate short read alignment with Burrows-Wheeler transform. Bioinforma. Oxf. Engl. 25, 1754–1760.

24. Li, H., Handsaker, B., Wysoker, A., Fennell, T., Ruan, J., Homer, N., Marth, G., Abecasis, G., Durbin, R. and 1000 Genome Project Data Processing Subgroup (2009). The Sequence Alignment/Map format and SAMtools. Bioinforma. Oxf. Engl. 25, 2078–2079.

25. Macdonald, R. L. and Gallagher, M. J. (2015). The Genetic Epilepsies. In Rosenberg’s Molecular and Genetic Basis of Neurological and Psychiatric Disease, pp. 973–998. Elsevier.

26. Macnamara, E. F., Schoch, K., Kelley, E. G., Fieg, E., Brokamp, E., Undiagnosed Diseases Network, Signer, R., LeBlanc, K., McConkie-Rosell, A. and Palmer, C. G. S. (2019). Cases from the Undiagnosed Diseases Network: The continued value of counseling skills in a new genomic era. J. Genet. Couns. 28, 194–201.

27. Maljevic Snezana, Krampfl Klaus, Cobilanschi Joana, Tilgen Nikola, Beyer Susanne, Weber Yvonne G., Schlesinger Friedrich, Ursu Daniel, Melzer Werner, Cossette Patrick, et al. (2006). A mutation in the GABAA receptor α1-subunit is associated with absence epilepsy. Ann. Neurol. 59, 983–987.

28. Monesson-Olson, B., McClain, J. J., Case, A. E., Dorman, H. E., Turkewitz, D. R., Steiner, B. and Downes, G. B. (2018). Expression of the eight GABAA receptor α subunits in the developing zebrafish central nervous system. PloS One 13, e0196083.

29. Nolan, D. and Fink, J. (2018). Chapter 30 - Genetics of epilepsy. In Handbook of Clinical Neurology (ed. Geschwind, D. H.), Paulson, H. L.), and Klein, C.), pp. 467–491. Elsevier.

30. Peng, X., Lin, J., Zhu, Y., Liu, X., Zhang, Y., Ji, Y., Yang, X., Zhang, Y., Guo, N. and Li, Q. (2016). Anxiety-related behavioral responses of pentylenetetrazole-treated zebrafish larvae to light-dark transitions. Pharmacol. Biochem. Behav. 145, 55–65.

31. Reyes-Nava, Nayeli G., Hernandez, Jose A., Castro, Victoria L., Reyes, Joel F., Castellanos, Barbara S. and Quintana, Anita M. (2018). Zebrafish: A Comprehensive Model to Understand the Mechanisms Underlying Neurodevelopmental Disorders. In Horizons in Neuroscience Research (ed. Costa, Andres. and Villalba, Eugenio., p. Nova Biomedical.

32. Samarut, É., Swaminathan, A., Riché, R., Liao, M., Hassan-Abdi, R., Renault, S., Allard, M., Dufour, L., Cossette, P., Soussi-Yanicostas, N., et al. (2018). γ-Aminobutyric acid receptor alpha 1 subunit loss of function causes genetic generalized epilepsy by impairing inhibitory network neurodevelopment. Epilepsia. 59, 2061–2074.

33. Sawyer, S. L., Hartley, T., Dyment, D. A., Beaulieu, C. L., Schwartzentruber, J., Smith, A., Bedford, H. M., Bernard, G., Bernier, F. P., Brais, B., et al. (2015). Utility of whole-exome sequencing for those near the end of the diagnostic odyssey: time to address gaps in care. Clin. Genet.

34. Sigel, E. and Steinmann, M. E. (2012). Structure, function, and modulation of GABA(A) receptors. J. Biol. Chem. 287, 40224–40231.

35. Tetreault, M., Bareke, E., Nadaf, J., Alirezaie, N. and Majewski, J. (2015). Whole-exome sequencing as a diagnostic tool: current challenges and future opportunities. Expert Rev. Mol. Diagn. 15, 749–760.

36. Thisse, C. and Thisse, B. (2008). High-resolution in situ hybridization to whole-mount zebrafish embryos. Nat. Protoc. 3, 59–69.

37. von Deimling, M., Häsler, R., Steinbach, V., Holterhus, P.-M., von Spiczak, S., Stephani, U., Helbig, I. and Muhle, H. (2017). Gene expression analysis in untreated absence epilepsy demonstrates an inconsistent pattern. Epilepsy Res. 132, 84–90.

38. Wangler, M. F., Yamamoto, S., Chao, H.-T., Posey, J. E., Westerfield, M., Postlethwait, J., Hieter, P., Boycott, K. M., Campeau, P. M. and Bellen, H. J. (2017). Model Organisms Facilitate Rare Disease Diagnosis and Therapeutic Research. Genetics. 207, 9–27.

39. Zhou, C., Ding, L., Deel, M. E., Ferrick, E. A., Emeson, R. B. and Gallagher, M. J. (2015). Altered Intrathalamic GABAA Neurotransmission in a Mouse Model of a Human Genetic Absence Epilepsy Syndrome. Neurobiol. Dis. 73, 407–417.

